# Semantic Segmentation of HeLa Cells: An Objective Comparison between one Traditional Algorithm and Three Deep-Learning Architectures

**DOI:** 10.1101/2020.03.05.978478

**Authors:** Cefa Karabağ, Martin L. Jones, Christopher J. Peddie, Anne E. Weston, Lucy M. Collinson, Constantino Carlos Reyes-Aldasoro

## Abstract

In this work, images of a HeLa cancer cell were semantically segmented with one traditional image-processing algorithm and three deep learning architectures: VGG16, ResNet18 and Inception-ResNet-v2. Three hundred slices, each 2000 × 2000 pixels, of a HeLa Cell were acquired with Serial Block Face Scanning Electron Microscopy. The deep learning architectures were pre-trained with ImageNet and then fine-tuned with transfer learning. The image-processing algorithm followed a pipeline of several traditional steps like edge detection, dilation and morphological operators. The algorithms were compared by measuring pixel-based segmentation accuracy and Jaccard index against a labelled ground truth. The results indicated a superior performance of the traditional algorithm (Accuracy = 99%, Jaccard = 93%) over the deep learning architectures: VGG16 (93%, 90%), ResNet18 (94%, 88%), Inception-ResNet-v2 (94%, 89%).

## 1 Introduction

The study of cells and their organelles have interested scientists from the early days of Hooke and van Leeuwenhoek to the formulation of cell theory by Schleiden and Schwann [1]. Since then, presence or absence of cells, shapes, inner components, interactions, regulation of processes, environment and many other characteristics have been thoroughly analysed, especially trying to relate these to conditions of health and disease [2–7]. To observe cells, it is necessary to use microscopy and one of its many different techniques like light, fluorescence or differential interference microscopy. Electron Microscopy (EM) can provide resolving power several orders of magnitude higher than conventional light and fluorescence microscopes and thus it is ideal to observe small structures of the cellular environment. Modern EM instruments allow the acquisition of contiguous images of a sample by slicing very thin sections from the top face of the resin-embedded sample with an ultramicrotome diamond knife [8]. Once the sample is sliced, the slice itself is discarded, the sample is raised into the imaging position and the scanning process continues for a given number of slices, thus creating a three-dimensional data set of contiguous images. This process is called *Serial blockface scanning EM* (SBF SEM) [9].

The nuclear envelope (NE) is a bi-layer membrane that separates the nucleus with the chromosomes from the rest of the cellular compartments [10] and contains a large number of membrane proteins with sophisticated roles and functions [11–14]. The structure and condition of the NE is of huge importance as it has been related to viral infections [15–19], Muscular dystrophy [20], Cancer [21–25], Osteoporosis [26], Cardiovascular diseases [27–29], other diseases [30–32], and ageing [33–35]. Therefore, algorithms for the segmentation, visualisation and analysis of the NE could provide parameters to understand the conditions of health and disease of a cell.

The segmentation and classification of cells and their environment through image-processing tasks have been important for many years and numerous algorithms have been proposed. Candia *et al.* summarised the importance of objective analysis emphatically in the following way: “we need unbiased, mathematically robust, scalable methods that allow us to identify key parameters that consistently characterize cell subpopulations … to build signatures of health and disease” [3]. PubMed [36] contains more than 33,000 entries with the words cell and classification or segmentation in the title and abstract ((classification[Title/Abstract] OR segmentation[Title/Abstract]) AND cell[Title/Abstract]). The number of entries drops considerably to less than 1000 when the keyword “electron” is added to the search. Segmentation and classification of images acquired with electron microscopy (EM) is difficult for several reasons. The considerable increase of size and resolution as compared with light and fluorescence microscopy provides complex morphological structures. Whilst fluorescence microscopy allows several channels that identify structures of interest, EM only provides a grey scale image and with a reduced contrast between the structures of interest and the background. Furthermore, when serial sections are obtained, the images are transformed into a volumetric data set.

Recently, advances in machine learning and artificial intelligence, especially those related to deep learning architectures [37], have revolutionised image processing tasks [38–43]. Several deep learning architectures [44–46] have obtained outstanding results in difficult tasks such as those of the ImageNet Large Scale Visual Recognition Challenge (ILSVRC) [47]. Not surprisingly, deep learning has become a popular tool for segmentation and classification. Convolution neural networks (CNN) [48], are versatile and have been shown to be very effective for a wide range of tasks including object detection [49, 50] image classification [51–55] and segmentation [56]. The U-Net architecture, proposed by Ronneberger [57] has become a widely used tool for segmentation and analysis. It recently became the most cited paper presented in the prestigious MICCAI conference. Cireşan [58] *et al.* applied deep neural networks (DNN) to detect membrane neuronal and mitosis detection in breast cancer [59]. Within EM studies, deep learning has been applied to analyse mitochondria [60, 61], synapses [62] and proteins [63].

Deep learning architectures have two main limitations: 1) they require a large amount of training data and 2) they require significant computational power. As graphics processing units (GPUs) become more popular, the main limitation is thus the scarcity of training data [52, 64–67].

In this work, three deep learning architectures, VGG16 [46], ResNet18 [68], and Inception-ResNet-v2 [69] were used to perform the semantic segmentation of HeLa cells. VGG16 has been widely used in a variety of image classification problems. ResNet solves the problem of vanishing/exploding gradients and was the winner of ILSVRC 2015 [47]. Inception-ResNet-v2 employs dropout to avoid overfitting and is seen as the successor of GoogLeNet [69]. These networks were selected due to their good balance between accuracy and computational complexity, especially ResNet and Inception-ResNet-v2, which outperform other common configurations and are at the Pareto frontier considering accuracy and complexity [70–72].

The algorithms were pre-trained with ImageNet and then fine-tuned with training data prepared for this work. These were then compared with a traditional image processing algorithm [73]. The image processing algorithm followed a pipeline of traditional tasks: low-pass filtering, edge detection, dilation, generation of super-pixels, distance transforms, mathematical morphology and post-processing to segment automatically the nuclear envelope and background of HeLa cells.

The results of the four algorithms were objectively compared against a ground truth that was formed by a manually segmented nuclear envelope and an automatically segmented background. All the programming was performed in Matlab^®^ (The Mathworks™, Natick, USA) and the codes are freely available through GitHub and the data sets through EMPIAR (see Supplementary Materials).

## 2 Materials and Methods

### 2.1 HeLa cells preparation and acquisition

Details of the cell preparation have been published previously [74], but briefly, the data set consisted of EM images of HeLa cells. HeLa cells were prepared and embedded in Durcupan resin following the method of the National Centre for Microscopy and Imaging Research (NCMIR) [75].

### 2.2 Image Acquisition

Once the cells were prepared, the samples were imaged using Serial Blockface Scanning Electron Microscopy (SBF SEM) with a 3View2XP (Gatan, Pleasanton, CA) attached to a Sigma VP SEM (Zeiss, Cambridge). The resolution of each image was 8, 192 × 8, 192 pixels corresponding to 10 × 10 nm (Fig. 1a). In total, the sample was sliced 517 times and corresponding images were obtained. The slice separation was 50 nm. The images were acquired with high-bit contrast (16 bit) and after contrast/histogram adjustment, the intensity levels were reduced to 8 bit and therefore the intensity range was [0 – 255]. Then, one cell was manually cropped by selecting its estimated centroid and a volume of 2, 000 × 2, 000 × 300 voxels was selected (Fig. 1b). Images are openly accessible via the EMPIAR public image database (http://dx.doi.org/10.6019/EMPIAR-10094).

**Fig 1.**
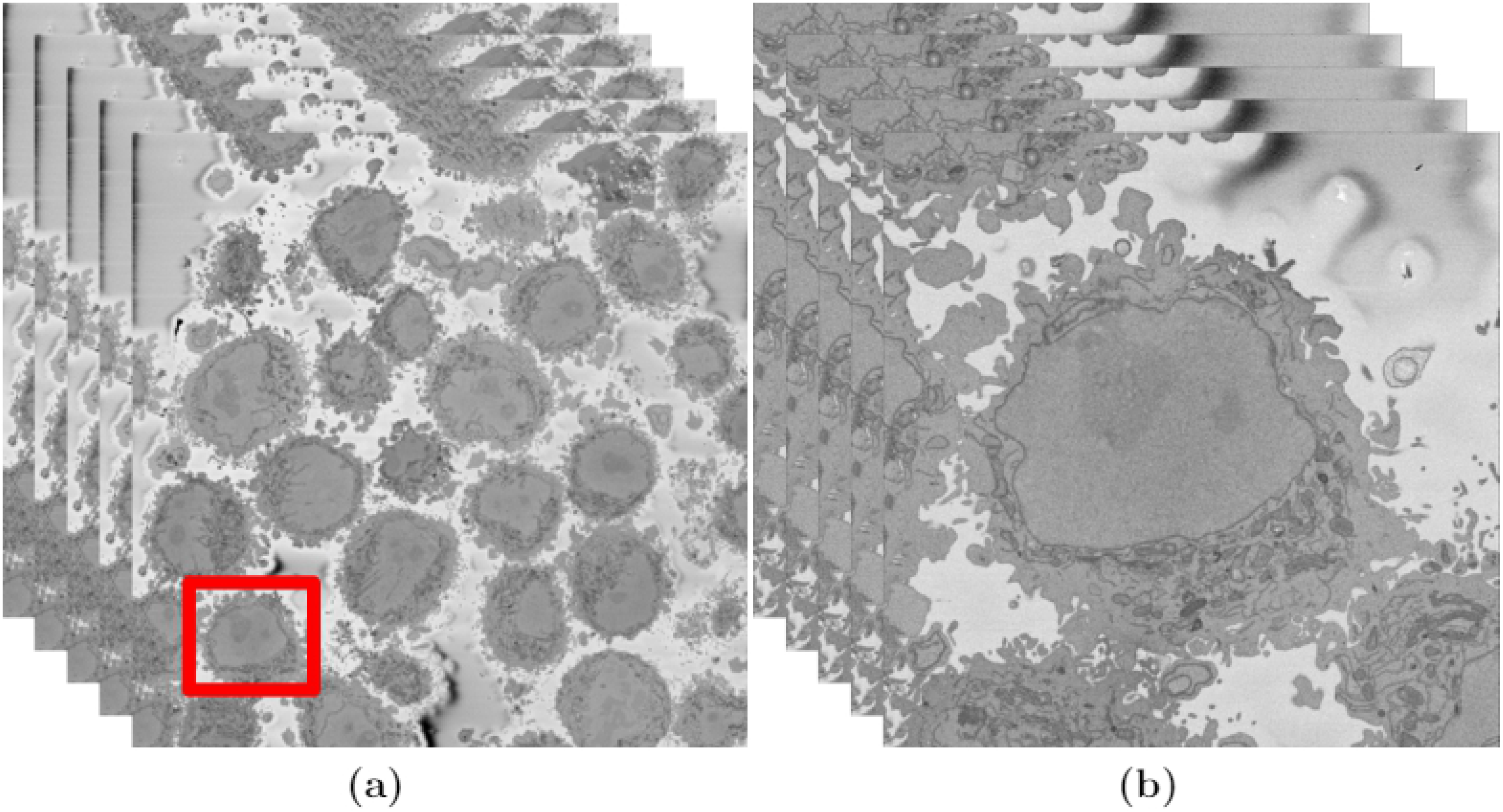
Illustration of the Serial Block Face Scanning Electron Microscope (SBF SEM) images containing HeLa cells. (a) Five representative 8192 × 8192 images arranged as 3D stack. The HeLa cells are the darker regions and the background is a brighter shade of grey. The red box indicates a region of interest (ROI), that is magnified on the right. (b) Detail of the ROI with a single cell in the centre. This is slice 118 of 300. The nucleus is the large and fairly uniform region in the centre and it is surrounded by the nuclear envelope (NE) which is darker than the nucleus.

### 2.3 Ground truth (GT) and training data

The three hundred slices were segmented with a combination of manual and algorithmic steps to provide a ground truth (GT). The NE was delineated manually using Amira (ThermoFisher Scientific, Waltham, MA, USA) and a Wacom (Kazo, Japan) Cintiq 24HD interactive pen display by one of the authors (A.E.W.) in approximately 30 hours (Fig. 2a). In order to determine whether disjoint regions belong to the nucleus, the user scrolled up and down through neighbouring slices to check connectivity of the regions. In a few cases, there were discontinuities in the line of the NE, and thus to morphological dilation was applied to ensure a closed contour.

**Fig 2.**
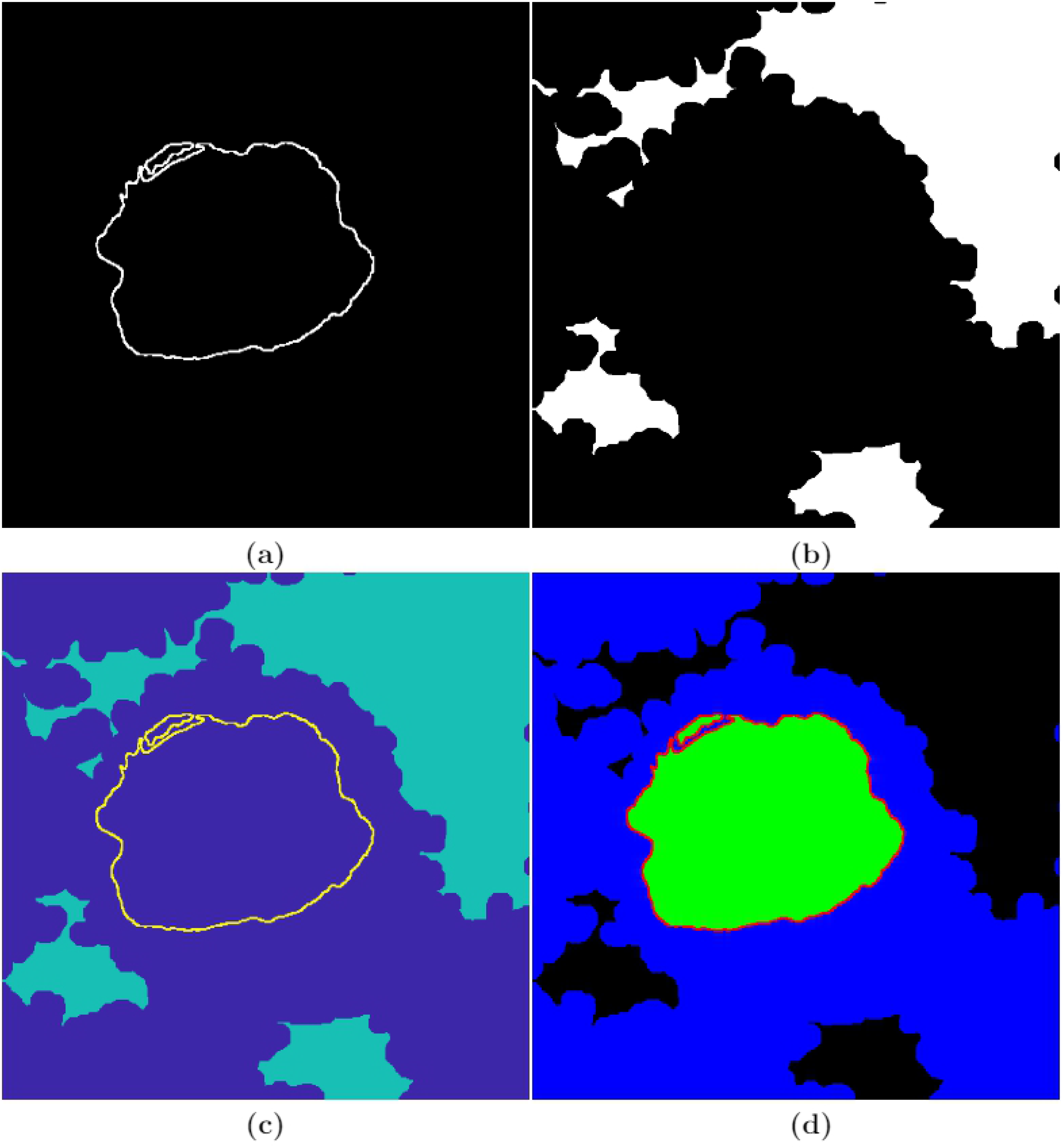
Illustration of the process followed to generate the ground truth. (a) Manually delineated nuclear envelope (NE). (b) Automatically segmented background. (c) Combination of the NE and the background. (d) Labelled image, generated in MATLAB^®^ *Image Labeler*, indicating the four different classes - nuclear envelope (red), nucleus (green), rest of the cell (blue), and background (black).

The background of a HeLa cell image was segmented automatically with an image-processing algorithm, which assumed that the background was brighter than the cells. The HeLa images were low-pass filtered with a Gaussian kernel with size *h* = 7 and standard deviation *σ* = 2 to remove high frequency noise. Canny edge detection was used to detect abrupt changes of intensity - edges. In order to to connect disjoint edges, they were further dilated. The complement of the edges (i.e. the regions where intensity was relatively uniform) were then labelled and its average intensity calculated. The background was selected as the brighter and larger regions previously segmented. Morphological operators were used to fill holes and close the regions for a more uniform background (Fig. 2b).

Next, the NE and the background were combined (Fig. 2c), exported to MATLAB^®^ *Image Labeler* with which four classes (nuclear envelope, nucleus, rest of the cell, background) were labelled (Fig. 2d). The GT was replicated to create an image with three channels to be consistent with the RGB images commonly used with pre-trained neural networks.

Ideally, the number of elements should be balanced between classes. However, the sizes of the classes in the HeLa data set were imbalanced, which is a common issue in biomedical imaging, especially the NE was relatively small as compared with the other classes. To improve training, class weighting was used to balance the classes. The pixel label counts computed earlier was used in order to calculate the median frequency class weights.

### 2.4 Semantic segmentation of HeLa cells

#### 2.4.1 Image-Processing Algorithm

The initial step of the image-processing algorithm [73] filtered the images with a low-pass filter with a Gaussian kernel with size *h* = 7 and standard deviation *σ* = 2 to remove high frequencies (Fig. 3a). The algorithm then exploited the abrupt change in intensity at the NE compared with the neighbouring cytoplasm and nucleoplasm by applying Canny edge detection [76]. To connect any disjoint edges, these were dilated by calculating a distance map from the edges and then all pixels within a certain distance were included as edges. The minimum distance was 5 and could grow according to the standard deviation of the Canny edge detector, which is an input parameter of the algorithm. These disjoint edges were part of the NE and were initially disjoint due to intensity variations in the envelope itself (Fig. 3b). The connected pixels not covered by the dilated edges were labelled by the standard 8-connected objects found in the image to create a series of superpixels (Fig. 3c). The superpixel size was not restricted so that large superpixels covered the background and nucleoplasm. Morphological operators were used to: remove regions in contact with the borders of the image, remove small regions, fill holes inside larger regions and close the jagged edges. From the volumetric perspective, the algorithm began at the central slice of the cell, which was assumed to be the one in which the nuclear region would be centrally positioned and have the largest diameter. The algorithm exploited the 3D nature of the data by propagating the segmented NE of a slice to the adjacent slices (up and down). The NE of a previous slice was used to check the connectivity of disjoint regions or *islands* separate from the main nuclear region. This is the same strategy that a human operator would follow by considering contiguous images and proceeded in both directions (up and down through the neighbouring slices) and propagated the region labelled as nucleus to decide if a disjoint nuclear region in the neighbouring slices (above or below) was connected above or below the current slice of analysis. When a segmented nuclear region overlapped with the previous nuclear segmentations, it was maintained, when there was no overlap, it was discarded (Fig. 3d). A different slice with disjoint areas is shown after image processing algorithm segmentation in Fig. 3e.

**Fig 3.**
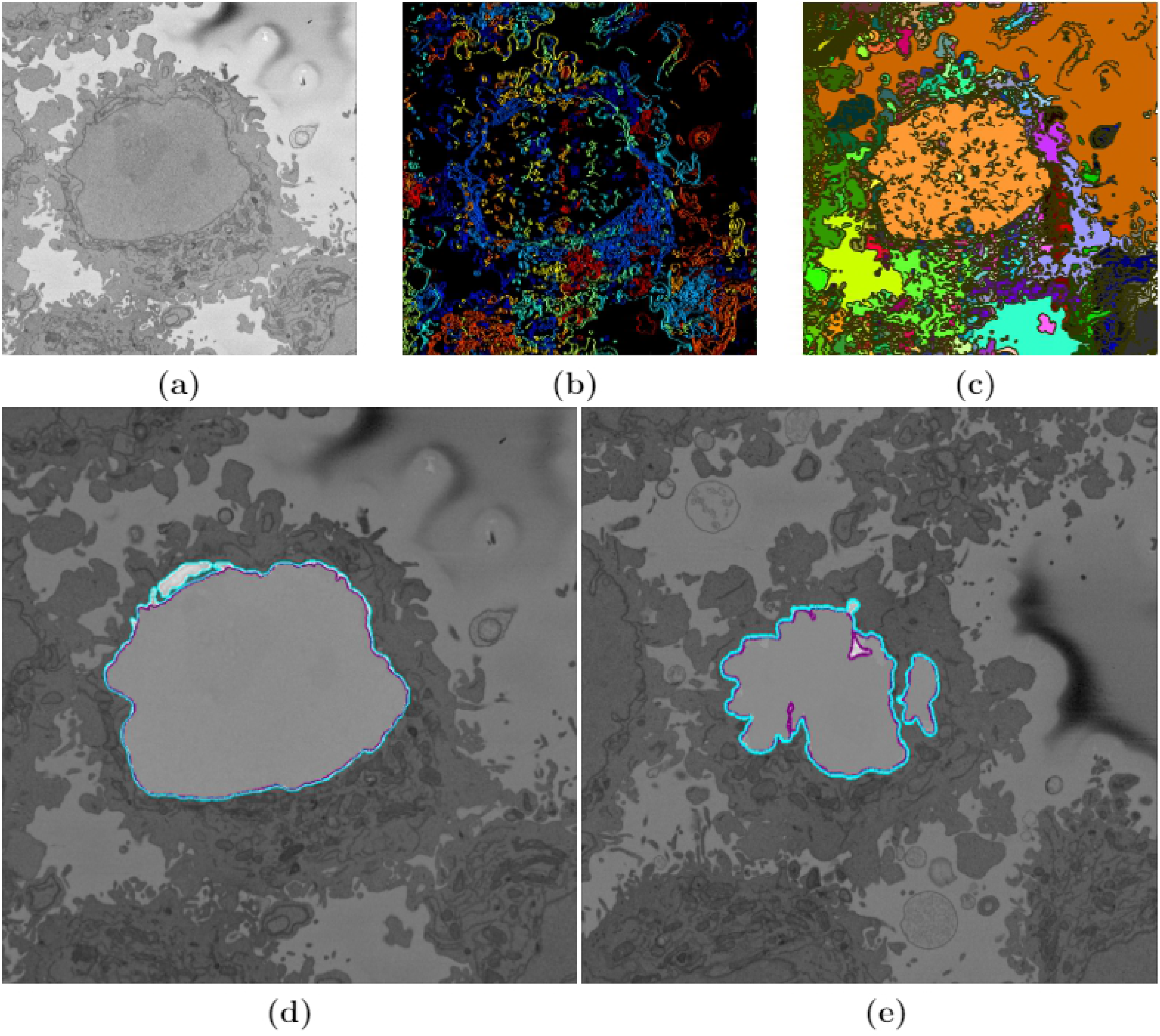
Illustration of intermediate steps of the proposed segmentation algorithm. (a) Cropped region around one HeLa cell (red box in Fig. 1a and Fig. 1b), surrounded by resin (background) and edges of other cells. This image was low-pass filtered. (b) Edges detected by Canny algorithm. The edges were further dilated to connect those edges that may belong to the nuclear envelope (NE) but were disjoint due to the variations of the intensity of the envelope itself. (c) Superpixels obtained with the image-processing algorithm and they were generated by removing dilated edges. Small superpixels and those in contact with the image boundary were discarded and the remaining superpixels were smoothed and filled, before discarding those by size that did not belong to the nucleus. (d) Final segmentation of the NE overlaid on the filtered image shown in purple. The manual segmentation or the ground truth (GT) is also shown in cyan. (e) A different slice showing final segmentation and GT overlaid on the filtered image. By using neighbouring segmentation as input parameter to the current segmentation and taking the regions into account, the segmentation was considerably improved and was able to identify disjoint regions as part of a single nucleus. Details of differences can be appreciated and the nuclear area covered by the GT and segmentation was brightened up for visualisation purposes.

#### 2.4.2 Deep learning architectures for semantic segmentation - VGG16, ResNet18 and Inception-ResNet-v2 Net configurations

A typical CNN combines a series of layers: convolutional layers followed by sub-sampling layers (Pooling layer), then another convolutional layers followed by pooling layers, and can continue for a certain number of times after which fully-connected layers are added to produce a prediction (e.g. estimated class probabilities). This layer-wise arrangement allows CNNs to combine low-level features to form higher-level features, learn features and eliminate the need for hand crafted feature extractors. In addition, the learned features are translation invariant, incorporate the two-dimensional (2D) spatial structure of images which contributed to CNNs achieving state-of-the-art results in image-related tasks [77].

The input to a CNN, i.e. an image to be classified, transits through the different layers to produce at the end some scores (one score per neuron in the last layer). In the case of image classification, these scores can be interpreted as the probability of the image to belong to each of the classes, which in this work are: nucleus, nuclear envelope, rest of the cell, and background. The goal of the training process is to learn the weights of the filters at the various layers of the CNN. The output of one of the layers before the last layer, which is fully connected, can be used as a global descriptor for the input image. The descriptor can then be used for various image analysis tasks including classification, recognition, and retrieval [48].

Three pre-trained deep neural networks, VGG16, ResNet18 and Inception-ResNet-v2 were fine-tuned to perform semantic segmentation of HeLa cells imaged with SBF SEM. These pre-trained deep neural networks have been widely explained in literature, but for completeness, a brief description of each architecture is given below.

##### VGG16

VGG16 is a convolution neural network (CNN) [46, 48], which takes as input 224 × 224 RGB images. The image is passed through a stack of convolutional layers, where filters are used with a very small receptive field: 3 × 3 (which is the smallest size to capture the notion of left/right, up/down, centre). In one of the configurations 1 × 1 convolution filters are utilised, which can be seen as a linear transformation of the input channels (followed by non-linearity). The convolution stride is fixed to 1 pixel; the spatial padding of convolution layer input is such that the spatial resolution is preserved after convolution, i.e. the padding is 1 pixel for 3 × 3 convolution layers. Spatial pooling is carried out by five max-pooling layers, which follow some of the convolution layers (not all the convolution layers are followed by max-pooling). Max-pooling is performed over a 2 × 2 pixel window, with stride 2. A stack of convolutional layers (which has a different depth in different architectures) is followed by three Fully-Connected (FC) layers: the first two have 4096 channels each, the third performs 1000-way ILSVRC classification and thus contains 1000 channels (one for each class). The final layer is the soft-max layer. The configuration of the fully connected layers is the same in all networks. All hidden layers are equipped with the rectification (ReLU [44]) non-linearity. The second column of Fig. 4 shows the basic network architecture of VGG16.

**Fig 4.**
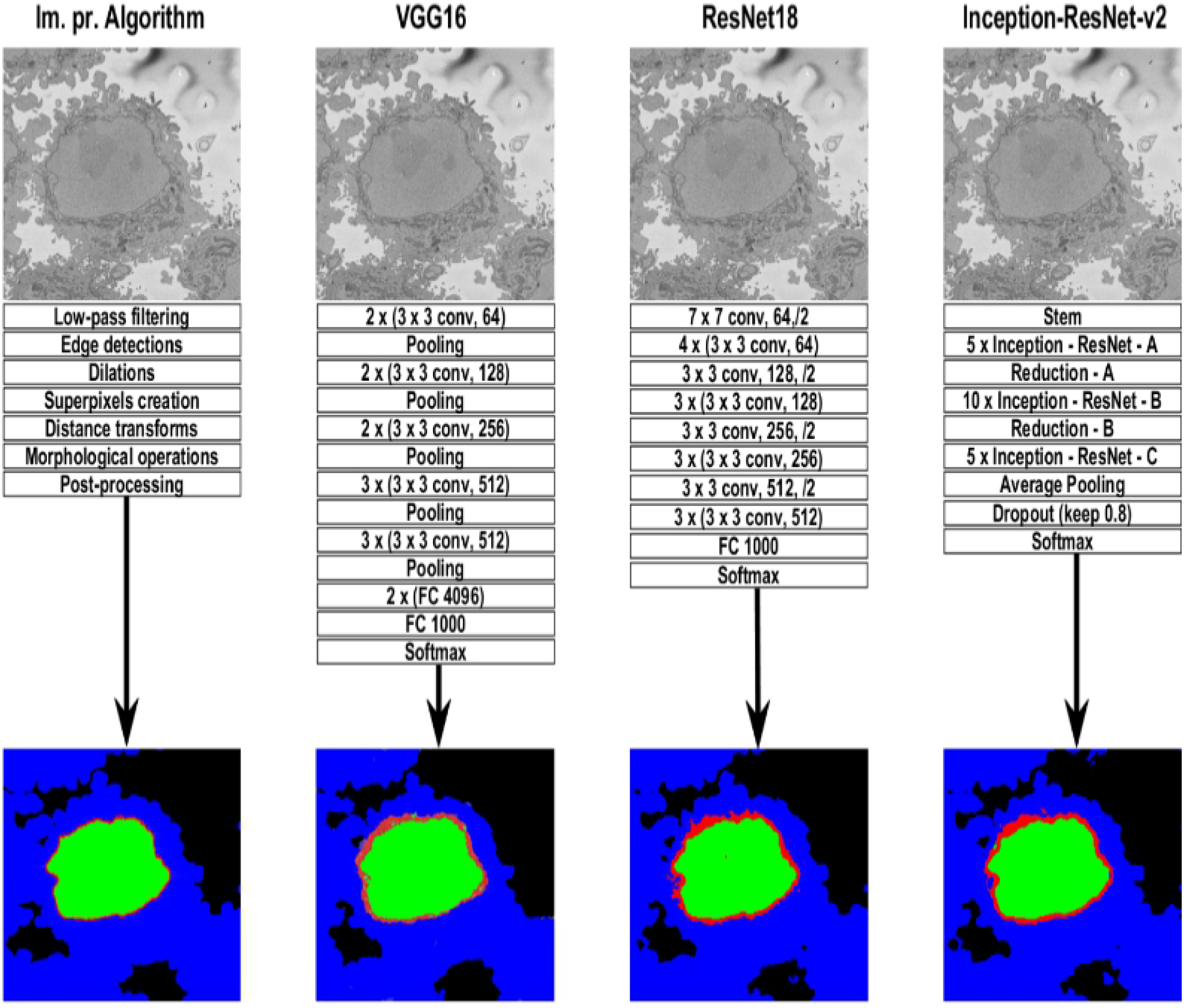
Graphical illustration of the four algorithms compared in this work. A representative slice of a HeLa cell image, the traditional image processing algorithm intermediate steps (first column), illustrations of three pre-trained deep learning architectures (Convolutional Neural Network (CNN)) - VGG16, ResNet18, and Inception-ResNet-v2 for the analysis of HeLa images, and semantic segmentation results showing four classes (nucleus, nuclear envelope, rest of the cell, and background) from all methodologies. A CNN is composed of an input layer, an output layer, and many hidden layers in between. These layers perform operations that alter the data with the intent of learning features specific to the data. Three of the most common layers are: convolution, activation or Rectified linear unit (ReLU), and pooling. An additional dropout layer is implemented in Inception-ResNet-v2 (last column) in order to avoid over-fitting. In this work, a stack of convolutional layers is followed by Fully-Connected (FC) layers and the softmax layer which performs a 4-way classification and thus contains 4 channels (one for each class - nucleus, nuclear envelope, rest of the cell, and background). These operations are repeated over tens or hundreds of layers with each layer learning to identify different features.

##### ResNet18

ResNet18 is mainly inspired by the philosophy of VGG16 [68], its total number of weighted layers is 18 and has an image input size of 224-by-224. The convolutional layers mostly have 3 × 3 filters and follow two simple design rules: (i) for the same output feature map size, the layers have the same number of filters; and (ii) if the feature map size is halved, the number of filters is doubled so as to preserve the time complexity per layer. Shortcut connections which turn the network into its counterpart residual version are inserted. The identity shortcuts can be directly used when the input and output are of the same dimensions. When the dimensions increase, two options are considered: (a) The shortcut still performs identity mapping, with extra zero entries padded for increasing dimensions. This option introduces no extra parameter; (b) The projection shortcut is used to match dimensions (done by 1 × 1 convolutions). For both options, when the shortcuts go across feature maps of two sizes, they are performed with a stride of 2. Down sampling is performed directly by convolutional layers that have a stride of 2. The network ends with a global average pooling layer and a 1000-way fully-connected layer with soft-max. ResNet18 has fewer filters and lower complexity than VGG16. The third column of Fig. 4 shows the basic network architecture of ResNet18.

##### Inception-ResNet-v2

The Inception deep convolutional architecture was introduced as GoogLeNet in [78] and named Inception-v1. Later the Inception architecture was refined in various ways, first by the introduction of batch normalisation [79] (Inception-v2). Later by additional factorisation ideas in the third iteration [80] which is referred to as Inception-v3. The Inception architecture is highly tunable, meaning that there are a lot of possible changes to the number of filters in the various layers that do not affect the quality of the fully trained network. The introduction of residual connections leads to dramatically improved training speed for the Inception architecture. For the residual versions of the Inception networks, Inception blocks in which a 5 × 5 convolution is replaced two 3 × 3 convolution operations to improve computational speed are used - stacking two 3 × 3 convolutions leads to a boost in performance. Each Inception block is followed by filter-expansion layer (1 × 1 convolution without activation) which is used for scaling up the dimensionality of the filter bank before the residual addition to match the depth of the input. This is needed to compensate for the dimensionality reduction induced by the Inception block.

Several versions of the residual version of Inception were tried. The first version, Inception-ResNet-v1, has roughly the computational cost of Inception-v3, while Inception-ResNet-v2 matches the raw cost of the newly introduced Inception-v4 network. However, the step time of Inception-v4 proved to be significantly slower in practice, probably due to the larger number of layers. The models Inception-v3 and Inception-v4 are deep convolutional networks not utilising residual connections while Inception-ResNet-v1 and Inception-ResNet-v2 are Inception style networks that utilise residual connections instead of filter concatenation.

Inception-ResNet-v2 is the combination of two of residual connections and the latest revised version of the Inception architecture [69] and it has an image input size of 299-by-299 and 164 layers deep. In the Inception-ResNet block, multiple sized convolutional filters are combined by residual connections. The usage of residual connections not only avoids the degradation problem caused by deep structures but also reduces the training time [81]. The 35 × 35, 17 × 17 and 8 × 8 grid modules, known as Inception-A, Inception-B and Inception-C blocks, are used in the Inception-ResNet-v2 network. The last column of Fig. 4 shows the basic network architecture of Inception-ResNet-v2.

Both ResNet18 and Inception-ResNet-v2 were fined tuned and trained on a new classification task - HeLa cells with four different classes. ResNet18 and Inception-ResNet-v2 are Directed Acyclic Graph (DAG) networks with branches which are faster, smaller and more accurate. ResNet18 has about 11.7 million (approx) parameters while Inception-ResNet-v2 has about 55.9 million (approx) parameters. ResNet18 and Inception-ResNet-v2 are convolutional neural networks that were trained on more than a million images from the ImageNet [82] database. They can classify images into 1000 object categories, such as keyboard, mouse, pencil, and many animals. As a result, the networks have learned rich feature representations for a wide range of images.

### 2.5 Description of Network Training

The HeLa Pixel-Labeled Images data set, shown in Fig. 2d, provides pixel-level labels for four semantic classes including nucleus, nuclear envelope, rest of the cell and background. These classes were specified before the training. The images and labelled training data in the HeLa data set are 2000 × 2000 × 1. In order to reduce training time and memory usage, all images and pixel label images were resized to 360 × 480 × 3. The network was trained using 60% of the images from the data set. The rest of the images (40%) were used to test the network after training. The network randomly splits the image and pixel label data into a training and test set. The whole data set for each pattern has been divided into two. Sixty percent is kept for training and its number has been increased artificially by using image augmentation techniques such as translation and reflections. EM images do not contain any colour information therefore they have the dimensions (*n_h_, n_w_, n_d_*) = (2000, 2000, 1). As all three deep neural networks (VGG16, ResNet18 and Inception-ResNet-v2) expect an image with *n_d_* = 3, the other 2 dimensions were a copy of the first dimension creating a greyscale image before training the algorithm.

The training took 21.25 hours on a single CPU and the training plot was obtained to check the accuracy and loss during training of the three pre-trained deep neural networks (VGG16, ResNet18 and Inception-ResNet-v2). The optimisation algorithm used for training is stochastic gradient descent with momentum (sgdm) and this was specified in training options. The sgdm algorithm can oscillate along the path of steepest descent towards the optimum. Adding a momentum term to the parameter update is one way to reduce this oscillation [83].

An image data augmenter in neural network configures a set of pre-processing options for image augmentation, such as resizing, rotation, and reflection and generates batches of augmented images. Data augmentation is used during training to provide more examples to the network because it helps improve the accuracy of the network, prevent the network from over fitting [54] and memorising the exact details of the training images. Three hundred images were used for training. These were split 60%/40% for training and testing. Data augmentation was applied with random translations of ±10 pixels on the horizontal and vertical axes and random reflection on the horizontal axis.

The training data and data augmentation selections were combined. The deep network reads batches of training data, applies data augmentation, and sends the augmented data to the training algorithm.

The VGG16 deep network training had 100 epochs with learning rate 0.001. Iterations per epoch was 45 therefore total number of iterations for the whole training was 4500. Similarly, training took approximately 1.5 hours for ResNet18 and 4.7 hours for Inception-ResNet-v2 with 30 epochs and 660 iterations and learning rate 0.00009 on a single CPU.

The semantic segmentation results from image-processing algorithm, VGG16, ResNet18, and Inception-ResNet-v2 are shown in Fig. 4. These results were compared with the labelled data shown in (Fig. 2d) and accuracy and Jaccard similarity index were calculated to asses the accuracy of the network. In order to measure accuracy for the data set, deep neural networks were run on the entire test set.

### 2.6 Quantitative Comparison

In order to evaluate the accuracy of the image processing segmentation algorithm and deep neural networks, two different pixel-based metrics were used: accuracy and Jaccard similarity index, or simply Jaccard index (JI) [84], were calculated. Both metrics arise from the allocation of classes to every pixel of an image, for which four cases exist: (i) true positive (TP), correspond to pixels which were correctly predicted as a certain class (e.g. nucleus) or to have a condition present (e.g. a disease), (ii) true negative, (TN) correspond to a pixel that was correctly predicted to be background or for which the condition not present (i.e. negative), (iii) false positive, (FP) correspond to those pixels predicted to be a class (or to have a condition) but correspond to a background (or not have the condition), and (iv) false negative (FN), correspond to those pixels that were predicted to be background (or not have the condition) but in reality belong to a class (or have the condition). Fig. 5 illustrates these cases for a samples slice of the data. Thus, accuracy can be defined mathematically in the following way:

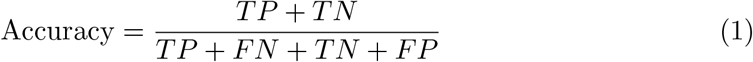

which corresponds to the sum of all correctly predicted pixels over the total number of pixels. Similarly, Jaccard index is calculated as:

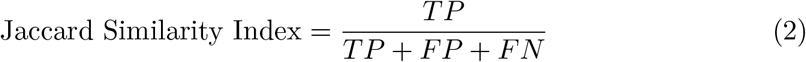

**Fig 5.**
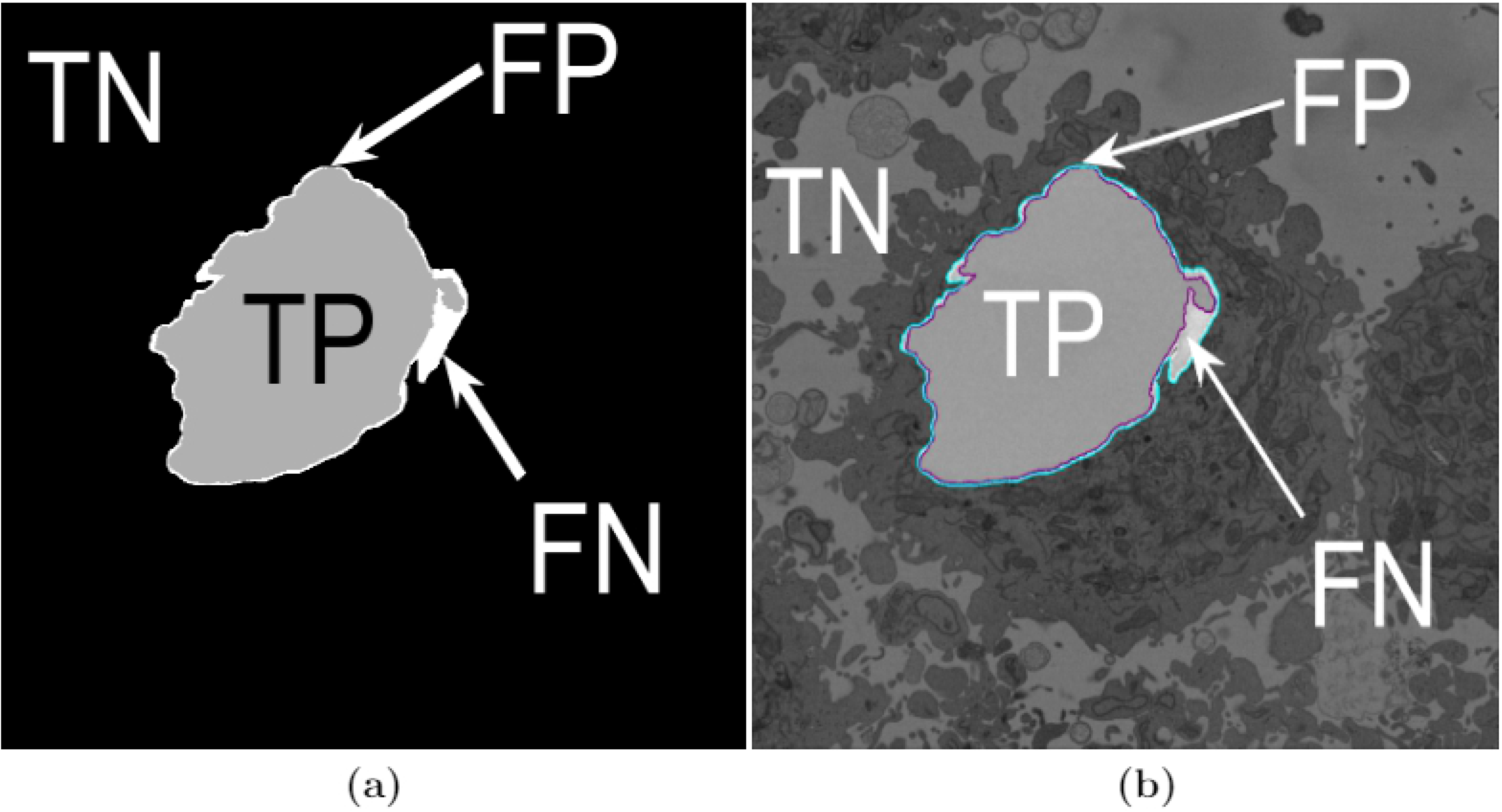
Illustration of the pixel-based metrics. (a) True Positives (TP, nuclear pixels segmented as nucleus), true negatives (TN, background pixels segmented as background), false positives (FP, background pixels segmented as nucleus) and false negatives (FN, nuclear pixels segmented as background). These quantities were used to compute accuracy and Jaccard similarity index for the image-processing algorithm and three pre-trained deep neural networks. (b) Segmentation overlaid on the same slice as (a) (slice 184/300 - on a filtered image): ground truth (GT) in cyan, automated segmentation in purple.

In both cases, the higher the number of the metric, the better the segmentation. It should be noticed that Jaccard is more rigorous as it does not take into account TN or background pixels, which in those cases where objects of interest are small in comparison with the image can bias a poor results to have a high accuracy. The metrics were calculated on a per-slice basis for all algorithms.

## 3 Results

In this work, images of HeLa cells observed with SBF SEM were semantically segmented with an image-processing algorithm and three pre-trained deep learning architectures (convolutional neural networks - CNNs), VGG16, ResNet18 and Inception-ResNet-v2. The accuracy and Jaccard similarity index of the segmentations against a ground truth were calculated. The image-processing algorithm is fully automatic and processed each slice of the HeLa cells in approximately 8 seconds and a whole cell containing 300 slices in approximately 40 minutes. On the other hand, the deep neural network architectures were fine-tuned and trained in 21.25 hours (VGG16), 1.5 hours (ResNet18), and 4.7 hours (Inception-ResNet-v2) to perform semantic segmentation on the whole cell with relatively good accuracy and differences can be appreciated in Fig. 6 with two ROIs.

**Fig 6.**
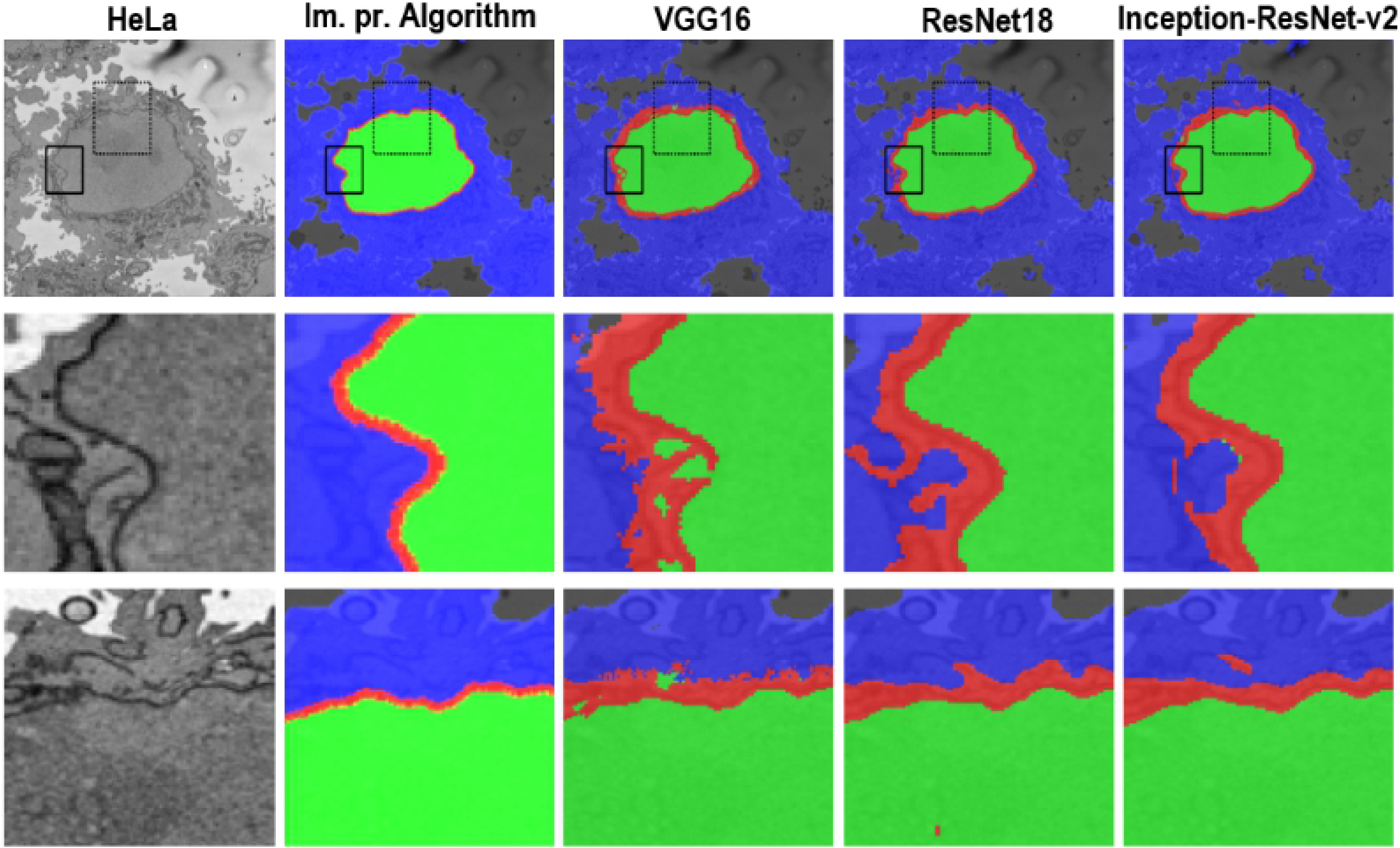
A representative HeLa image and semantic segmentation results overlaid on the filtered image. Top row: Boxes denote two Region of Interests (ROIs) on filtered image as well as semantic segmentation results. Results were obtained from all algorithms - a traditional and three deep learning architectures. The ROIs are magnified and shown in the middle and bottom rows. Middle row: ROI 1 corresponding black box with solid line. Bottom row: ROI 2 - box with dashed line. Differences in segmentation results can be seen from these ROIs - i.e. the red dot obtained from ResNet18 semantic segmentation on bottom row and 4^*th*^ column.

Segmentation of the nuclear envelope by all three deep neural networks was outperformed by the image-processing algorithm as shown in Fig. 6 and Fig. 7. Visually, the semantic segmentation results overlap well for classes such as nucleus, rest of the cell, and background. However, smaller objects like the nuclear envelope are not as accurate. Although the overall data set performance is quite high, the class metrics show that under represented classes such as nuclear envelope is not segmented as well as classes such as nucleus, rest of the cell, and background.

**Fig 7.**
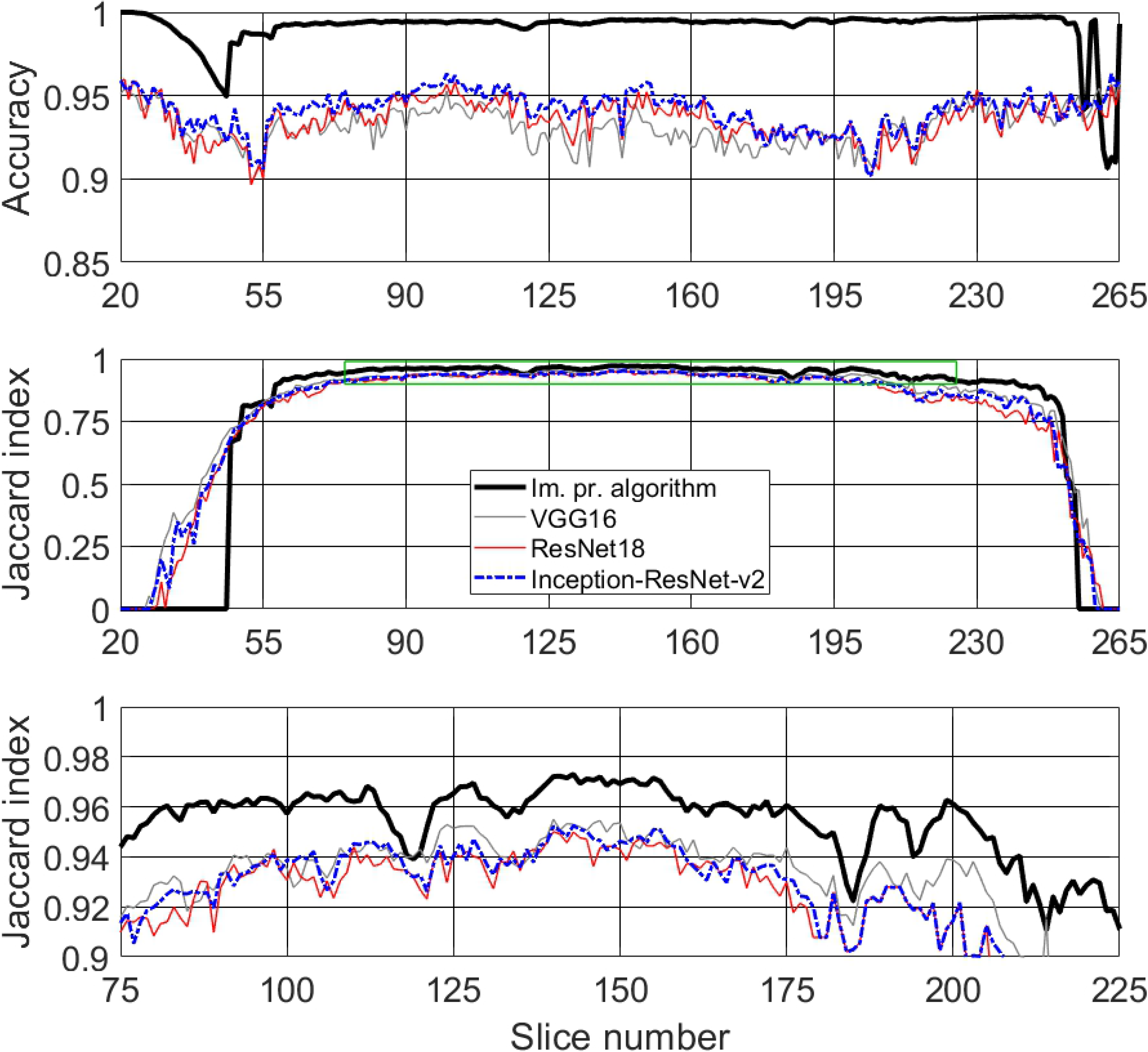
Metrics used for comparison between image-processing algorithm and three pre-trained deep neural networks - VGG16, ResNet18 and Inception-ResNet-v2 for semantic segmentation of HeLa cell imaged with electron microscopy (EM). (Top row) Accuracy. (Middle row) Jaccard similarity index, also known as intersection over union, for all algorithms. Green box denotes the central slices and corresponding Jaccard similarity index that is magnified below. (Bottom row) Jaccard similarity index for central slices (slices between 75/300 and 225/300 - interquartile range (IQR)), for easier comparison. The image-processing algorithm outperforms all deep neural networks.

## 4 Discussion

In this paper, a classical and unsupervised image processing algorithm was used to perform semantic segmentation of cancerous HeLa cell images from SBF SEM and compared with three pre-trained deep neural network architectures. The pre-trained deep neural network architectures, VGG16, ResNet18 and Inception-ResNet-v2 were trained in ImageNet and fine-tuned for semantic segmentation of the HeLa cells. Four different classes, nucleus, nuclear envelope, rest of the cell, and background were used in labelling training data for deep neural networks. Two similarity metrics, accuracy and Jaccard index, were calculated so the image-processing algorithm was compared with deep neural networks. For central slices, slices between 75/300 and 225/300, the image-processing algorithm outperformed all three deep neural networks as proved by the similarity metrics. As the image-processing algorithm was trained and tested only for the nuclear envelope and nucleus and as the bottom 26 and the top 40 slices did not contain any cell, the difference in Jaccard index, which can be observed away from central slices, is in favour of the deep neural networks towards the top and bottom of the cells (Fig. 7). Additional data that includes more samples of the under represented classes, nuclear envelope in this case, might help improve the results as the sizes of the classes in the HeLa data set were imbalanced. For the image-processing algorithm this would be expected, as the NE is more irregular on the top and bottom slices than on the central ones. The segmentation of each cropped cell is fully automatic and unsupervised and the image-processing algorithm segments one slice in approximately 8 seconds and one whole cell in approximately 40 minutes with good accuracy.

As JI does not count true negative (TN), the values decrease towards the top and bottom slices of the cells as the structure was considerably more complex and the areas become much smaller (Fig. 7 (Middle row) and (Bottom row)). On the other hand, accuracy includes the TN in both numerator and denominator and this, especially in cases where the objects of interest are small and there are large areas of background (e.g. the top and bottom slices of the cell) would render very high accuracy. Therefore, in contrast to JI, accuracy increases in slices towards both top and bottom ends.

Overall, the best results were obtained by the image processing segmentation algorithm especially for central slices - slices between 75/300 and 225/300 (Fig. 7 (Bottom row)).

Deploying deep learning architectures [37] and training to learn patterns and features directly from EM images and to automatically segment the NE and other parts of a cell is indeed necessary as this will reduce the effort, assessment variability and provide a second opinion to support biomedical researchers’ decisions as it shortens the time required to segment the cell.

The main contributions of this work are: (a) Generating 300 labelled images of the HeLa cell, shown in (Fig. 1), defined by four different classes - nuclear envelope, nucleus, rest of the cell, and background. These images were used to train deep learning architectures. (b) an objective comparison, supported by accuracy and Jaccard similarity index, between the image-processing algorithm and three pre-trained deep neural networks - VGG16, ResNet18 and Inception-ResNet-v2 to perform semantic segmentation of HeLa cells defined by four different classes - nuclear envelope, nucleus, rest of the cell, and background.

The open-source image-processing algorithm provides an alternative to expensive commercial software and manual segmentation, which is still widely used despite the significant disadvantages of time and inter- and intra-user variability.

## Supplementary Materials

The codes associated with this article are available open source at https://github.com/reyesaldasoro/Hela-Cell-Segmentation, https://github.com/karabagcefa/Hela-Cell-Semantic-Segmentation, and the data at http://dx.doi.org/10.6019/EMPIAR-10094 respectively.

## Author Contributions

M.L.J., C.J.P., A.E.W., and L.M.C. prepared the cells, imaged the data and segmented the ground truth of one cell, C.K. generated labelled images of the same cell in MATLAB^®^ *Image Labeler* so they could be used for training of the three pre-trained deep neural networks. C.K. and C.C.R-A. conceived, designed and performed the computational experiments; C.K. and C.C.R-A. analysed the data; C.K. and C.C.R-A. wrote the manuscript. M.L.J., A.E.W, and L.M.C. revised the manuscript. All authors approved the final submission paper.

## Acknowledgements

This work was supported by the *Francis Crick Institute*. The authors acknowledge the support of the *Alan Turing Institute* through the Data Study Groups organised by Dr Sebastian Vollmer where initial study of this data was made.

